# Comparative genomics on cultivated and uncultivated, freshwater and marine *Candidatus* Manganitrophaceae species implies their worldwide reach in manganese chemolithoautotrophy

**DOI:** 10.1101/2021.11.15.468770

**Authors:** Hang Yu, Grayson L. Chadwick, Usha F. Lingappa, Jared R. Leadbetter

**Affiliations:** Division of Geological & Planetary Sciences, California Institute of Technology, Pasadena, CA, USA 91125; Division of Engineering & Applied Science, California Institute of Technology, Pasadena, CA, USA 91125

**Keywords:** autotroph, lithotroph, chemolithoautotroph, manganese oxide, manganese carbonate, Nitrospirae, Nitrospirota, Mn, Mn2+, Mn(II), Mn (II)

## Abstract

Chemolithoautotrophic manganese oxidation has long been theorized, but only recently demonstrated in a bacterial co-culture. The majority member of the co-culture, *Candidatus* Manganitrophus noduliformans, is a distinct but not yet isolated lineage in the phylum *Nitrospirota* (*Nitrospirae*). Here, we established two additional MnCO_3_-oxidizing cultures using inocula from Santa Barbara (USA) and Boetsap (South Africa). Both cultures were dominated by strains of a new species, designated *Candidatus* Manganitrophus morganii. The next abundant members differed in the available cultures, suggesting that while *Ca*. Manganitrophus species have not been isolated in pure culture, they may not require a specific syntrophic relationship with another species. Phylogeny of cultivated *Ca*. Manganitrophus and related metagenome-assembled genomes revealed a coherent taxonomic family, *Candidatus* Manganitrophaceae, from both freshwater and marine environments and distributed globally. Comparative genomic analyses support this family being Mn(II)-oxidizing chemolithoautotrophs. Among the 895 shared genes were a subset of those hypothesized for Mn(II) oxidation (Cyc2 and PCC_1) and oxygen reduction (TO_1 and TO_2) that could facilitate Mn(II) lithotrophy. An unusual, plausibly reverse Complex 1 containing 2 additional pumping subunits was also shared by the family, as were genes for the reverse TCA carbon fixation cycle, which could enable Mn(II) autotrophy. All members of the family lacked genes for nitrification found in *Nitrospira* species. The results suggest that *Ca*. Manganitrophaceae share a core set of candidate genes for the newly discovered manganese dependent chemolithoautotrophic lifestyle, and likely have a broad, global distribution.

**Importance:** Manganese (Mn) is an abundant redox-active metal that cycled in many of Earth’s biomes. While diverse bacteria and archaea have been demonstrated to respire Mn(III/IV), only recently have bacteria been implicated in Mn(II) oxidation dependent growth. Here, two new Mn(II)-oxidizing enrichment cultures originated from two continents and hemispheres were examined. By comparing the community composition of the enrichments and performing phylogenomic analysis on the abundant *Nitrospirota* therein, new insights are gleaned on cell interactions, taxonomy, and machineries that may underlie Mn(II)-based lithotrophy and autotrophy.

## Introduction

Members of the bacterial phylum *Nitrospirota* (formerly *Nitrospirae*) are best known for having physiologies that exploit the utilization of high potential electron donors or low potential electron acceptors (1, 2). Cultivated organisms representing this phylum cluster within 4 clades. Order *Nitrospirales* (formerly genus *Nitrospira*) plays an important role in the nitrogen cycle, carrying out nitrite oxidation (3, 4) and complete ammonium oxidation to nitrate (5, 6). Class *Leptospirilla* (formerly genus *Leptospirillum*) thrive in low pH environments oxidizing iron (7). Class *Thermodesulfovibria* (formerly genus *Thermodesulfovibrio*) includes high temperature dissimilatory sulfate-reducers (8), some with the capacity of S disproportionation (9), as well as uncultivated magnetotactic bacteria (10). Recently, a bacterial co-culture was demonstrated to perform Mn(II) oxidation dependent chemolithoautotrophic growth (11). This metabolism was attributed to a member of a previously uncultivated clade of *Nitrospirota, Candidatus* Manganitrophus noduliformans strain Mn1, given that the minority member in the co-culture, *Ramlibacter lithotrophicus* (*Comamonadaceae*; formerly within the *Betaproteobacteria*, now within *Gammaproteobacteria*) could be isolated yet would not oxidize Mn(II) alone (11). Based on 16S rRNA gene phylogeny, several relatives of strain Mn1 were identified (11). However, whether or not these relatives might share the same Mn(II) oxidation metabolism was not something that could be gleaned from their rRNA genes.

Mn is the third most abundant redox-active metal in the Earth’s crust and is actively cycled (12– 14). Microbial reduction of Mn oxides for growth has been demonstrated in numerous bacterial and archaeal phyla (14–18). The notion that microbial oxidation of Mn(II) with O_2_ could serve as the basis for chemolithoautotrophic growth was first theorized decades ago (13, 14, 19, 20). This metabolism, while energetically favorable (ΔG°′ = −68 kJ/mol Mn), poses a biochemical challenge to the cell because of the high average potential of the two Mn(II)-derived electrons (Mn(II)/Mn(IV), E°′ = +466 mV (11)). These electrons would need their redox potential to be lowered by nearly a full volt in order to reduce the ferredoxin (E°′ = −320 to -398 mV (21)) employed in their CO_2_ fixation pathway (11). This is a larger and more significant mismatch in redox potential than similar chemolithotrophic metabolisms, such as nitrite or iron oxidation (NO_2^-^_ /NO_3^-^_, E°′ = +433 mV (21); Fe(II)/Fe(III), E°′ ∼ 0 mV (22)). Based on deduced homology with characterized proteins involved with Fe(II) oxidation or aerobic metabolism, genes for 4 putative Mn-oxidizing complexes and 5 terminal oxidases were identified in strain Mn1 and proposed as candidates for energy conservation via electron transport phosphorylation (11). Remarkably, gene clusters for 3 different Complex I exist in strain Mn1 and could facilitate the otherwise endergonic coupling of Mn(II) oxidation to CO_2_ reduction, allowing for autotrophic growth via reverse electron transport, i.e. expending motive force to drive down electron reduction potential (11). The apparent redundancy of diverse novel complexes in several members of the family remains puzzling. It seems clear that the identification and analysis of additional strains and genomes of Mn(II)-oxidizing chemolithoautotrophs could likely shed light on the complexes essential for this newfound mode of metabolism.

The ever increasing number of metagenome-assembled genomes (MAGs) available in the databases provides for an unprecedented opportunity to learn about the gene content and potential functions of many uncultured microorganisms. Yet, cultivation remains critical to forming interconnections between the genomes of both cultured and uncultivated microbes and their metabolisms. Herein, we successfully established new in vitro enrichment cultures performing chemolithoautotrophic Mn oxidation from two disparate environmental inoculum sources. By comparing the MAGs of the most abundant organisms present in these enrichments, also members of the *Nitrospirota*, as well as 66 newly and publicly available MAGs in the databases belonging to Nitrospirota clades with unexamined metabolisms, we gain insight into a core set of candidate genes for facilitating chemolithoautotrophic Mn oxidation, as well as the phylogenetic and geographic distribution of known and putatively Mn-oxidizing *Nitrospirota*.

## Results

### Reproducible cultivation of Mn-oxidizing chemolithoautotrophs

*Ca*. Manganitrophus noduliformans strain Mn1 was accidentally enriched in tap water (11). Using the defined Mn(II) carbonate medium in this previous study (11), new Mn-oxidizing enrichment cultures were successfully established from two distinct sample sources. One inoculum was material from a Mn oxide containing rock surface near Boetsap, Northern Cape, South Africa (“South Africa enrichment”), and the other inoculum was material from an iron oxide microbial mat in Santa Barbara, California, USA (“Santa Barbara enrichment”). While the new enrichments grew in the same defined freshwater medium, they exhibited different temperature optima. The South Africa enrichments initially grew at 28.5°C, although they oxidized Mn(II) faster at 32°C, similar to the previous enrichment from the Pasadena drinking water distribution system (“Pasadena enrichment”) (11). The Santa Barbara enrichments grew at 28.5°C, but not at 32°C. Otherwise, no striking differences in appearance (e.g. formation of small Mn oxide nodular products) between the three cultures was observed. These results indicate that the defined Mn(II) carbonate medium can successfully be employed during intentional, directed attempts to cultivate Mn-oxidizing chemolithoautotrophs from diverse terrestrial and aquatic freshwater environments.

### Community analysis of Mn-oxidizing enrichment cultures from three origins

As was the case with cultures of *Ca*. M. noduliformans, repeated attempts to identify single colonies of the lithotrophs responsible for Mn oxidation were not successful on an agar-solidified, defined Mn(II) carbonate medium. Sequencing of partial 16S rRNA genes amplified from the liquid cultures revealed differences in community structures between the Mn-oxidizing enrichments. The most abundant microorganism from the South Africa and Santa Barbara enrichments belonged to the same taxon as the previously described *Ca*. M. noduliformans (Figure 1). However, the identities of the next most abundant members of the communities differed. The previously described Pasadena enrichment containing *Ca*. M. noduliformans had *Ramlibacter lithotrophicus* as the second most abundant member throughout the enrichment refining process (Supplementary Table 1). *R. lithotrophicus* could be isolated from the enrichment using the same defined medium but with other electron donors such as succinate and hydrogen, but could not oxidize Mn(II) as an isolate (11). Organisms belonging to the same taxon as *R. lithotrophicus* were present in the South Africa enrichments, varying from 2-28 in rank abundance, but were not abundant in Santa Barbara enrichments (<0.5% relative abundance) (Figure 1 and Supplementary Table 1). In the South Africa enrichments, the second most abundant member varied between a *Pseudomonas* species (*Gammaproteobacteria*), a member of the *Zavarziniales* (*Alphaproteobacteria*), *R. lithotrophicus*, and *Hydrogenophaga* (a *Comamonadaceae* closely related to *R. lithotrophicus*) (Figure 1). In the Santa Barbara enrichments, the second most abundant member was a member of the *Anaerolineaceae* (phylum *Chloroflexi* or *Chloroflexota*; Figure 1). Changing the incubation temperature did not affect the identities of the 3 most abundant taxa in the South Africa enrichments (Figure 1). However, the choice of nitrogen source in the medium resulted in a shift in community member relative abundances (Figure 1). Notably, the only other shared organism between South Africa, Santa Barbara and Pasadena enrichments with >1% relative abundance was a member of the *Zavarziniales* (Figure 1 and Supplementary Table 1). Its relative abundance markedly increased when the South Africa enrichments were grown in medium with nitrate instead of ammonia as the nitrogen source. Overall, while the community composition varied between the Mn-oxidizing enrichments, strains of *Ca*. Manganitrophus were consistently the most abundant species in all such cultures.

**Figure 1.**
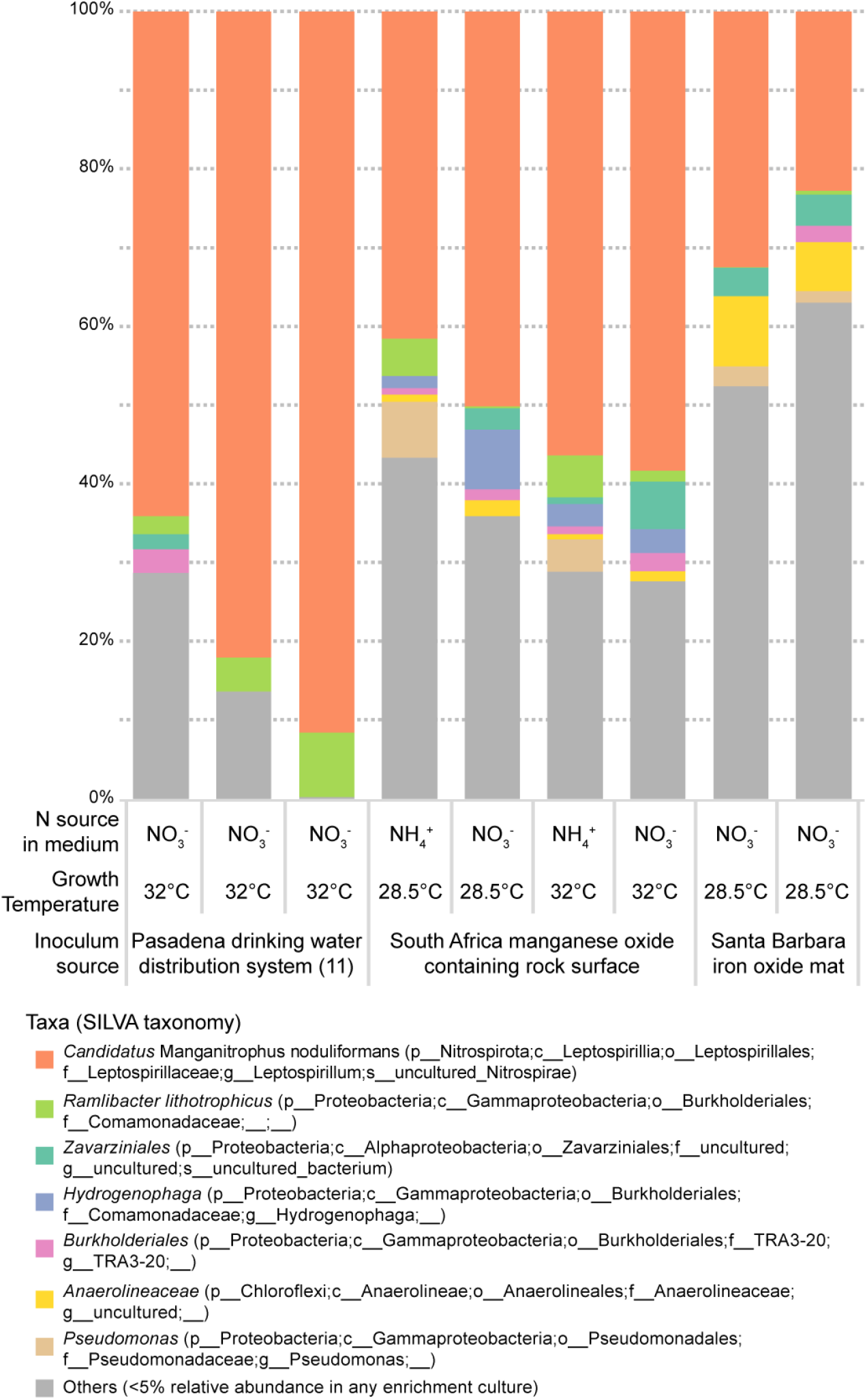
Community analysis of manganese-oxidizing enrichment cultures using partial 16S rRNA gene amplicon sequencing. Taxonomic classification is based on the SILVA SSU rRNA database v138. Detailed taxon relative abundances can be found in Supplementary Table 1.

### Expansion of metagenome-assembled genomes of cultivated and environmental Mnoxidizing *Nitrospirota*

We performed shotgun metagenomic sequencing on two of the new Mn-oxidizing enrichments in order to gain phylogenetic and functional insights into the newly cultivated *Ca*. Manganitrophus strains. We reconstructed high-quality MAGs (>97% completeness, <5% contamination) (23) of the most abundant organism from each metagenome (Supplementary Table 1). We refer to these MAGs as strain SA1 and SB1 to indicate that they originated from South Africa and Santa Barbara, respectively. Both genome and 16S rRNA gene phylogenies confirmed that strain SA1 and strain SB1 were related to the previously characterized *Ca*. M. noduliformans strain Mn1 (Figure 2). Based on their average nucleotide identities (ANI) and using 95% ANI as a possible metric for species delineation (24–26), strains SA1 and SB1 were provisionally considered to represent distinct strains of the same species (96% ANI). Both could be considered a different species than strain Mn1 (94% ANI) (Supplementary Table 3). The genome sizes of these 2 new strains were smaller (4.3 Mb) than that of strain Mn1 (5.2 Mb) (Supplementary Table 2). The arrangement of homologous regions in strains SA1 and SB1 were similar (Supplementary Figure 1a), but were different from strain Mn1 (Supplementary Figure 1b).

**Figure 2.**
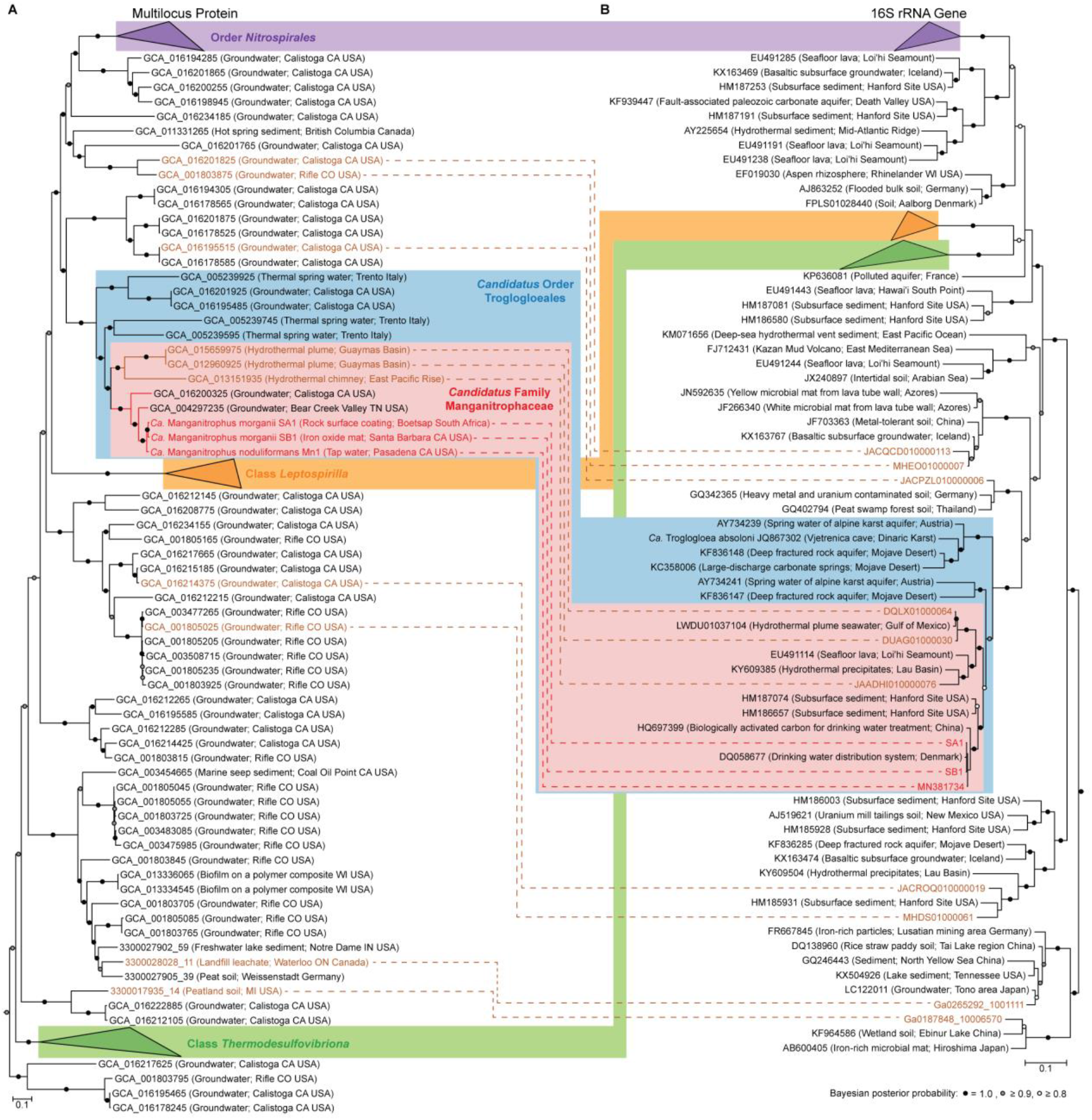
Phylogenetic analysis of the bacterial phylum Nitrospirota. **A**, Multilocus phylogram, based on a Bayesian analysis of 5040 aligned amino acid positions concatenated from 120 bacterial protein markers. **B**, 16S rRNA gene phylogram, based on a Bayesian analysis of 1508 aligned nucleotide positions. For both **A** and **B**, NCBI accession numbers or IMG contigs identifiers for the genome assemblies or 16S sequences are in the node names, with their source environments shown in brackets. Two phylograms can be linked by the genomes assemblies that contain 16S rRNA genes, with environmental metagenomes in brown and manganese-oxidizing enrichment cultures in red. Previously described taxonomic groups based on GTDB taxonomic classifications and the proposed taxonomic groups are color grouped.

These differences were also observed at the deduced protein level, with strains SA1 and SB1 more closely related to each other than to strain Mn1 (Supplementary Table 4). These variations in the proteins were not concentrated in one genomic region, but instead scattered throughout the genome (Supplementary Figure 1c). Further, de novo gene clustering showed that strains SA1 and SB1 shared more genes with each other than with strain Mn1 (Supplementary Figure 1d). All together, our results support strains SA1 and SB1 as a distinct species, which we designate as *Candidatus* Manganitrophus morganii (Supplementary Text). These 3 cultivated *Ca*. Manganitrophus strains in two different species provide a basis to examine the phylogenetic and genomic diversity of their shared metabolism, namely Mn-oxidizing chemolithoautotrophy.

In addition to reconstructing MAGs from Mn-oxidizing enrichments, we also analyzed publicly available MAGs in the phylum *Nitrospirota*. We screened for MAGs that did not belong in the three characterized clades, namely *Nitrospirales, Leptospirilla* and *Thermodesulfovibria*. As of 26 March 2019, only 3 MAGs had met this taxonomic criteria with completeness >50% and contamination <5% (11). However, as of March 30 2021, 64 new public high-quality (>90% completeness, <5% contamination) and 2 medium-quality (>50% completeness, <10% contamination) MAGs meeting this taxonomic criteria had become available (Supplementary Table 5). These 66 MAGs allowed for a much more detailed phylogenomic view into the uncultivated *Nitrospirota* and their potential ability to oxidize Mn.

### 16S rRNA gene and multilocus protein phylogeny reveal robust taxonomic groups

The available MAGs provide a phylogenetic resolution that matches the traditionally employed 16S rRNA genes (Figure 2). The MAGs were spread out across different phylogenetic clusters within the phylum (Figure 2a). Using the 14 MAGs that also contained 16S rRNA genes, we were able to link the genome phylogeny to the 16S rRNA gene phylogeny, and observed similar clusterings between the two phylogenetic approaches (Figure 2). The 3 cultivated strains all resided within the genus *Ca*. Manganitrophus. Other members of *Ca*. Manganitrophus, based on either their genomes or 16S rRNA genes, were from terrestrial, aquatic and engineered environments, and all freshwater in origin (Figure 2). Our phylogeny revealed a sister genus of marine origin (Figure 2). Together, these two genera form a coherent and well supported phylogenetic clade, hereafter termed family *Candidatus* Manganitrophaceae (Figure 2).

Previously, the class *Candidatus* Troglogloea was proposed to encompass strain Mn1 and *Candidatus* Troglogloea absoloni (an uncultivated species from Vjetrenica cave in the Dinaric Karst), based on their 16S rRNA gene phylogeny (11). Based on our new phylogenomic analysis, we propose that the order *Ca*. Troglogloeales includes the family *Ca*. Manganitrophaceae, *Ca*. T. absoloni, and its relatives (Figure 2), together constituting a sister group distinct from the order *Nitrospirales* (which includes the cultivated nitrite and ammonia-oxidizing *Nitrospirota*). These genera, family, and order proposals are consistent with the latest taxonomic classification in the Genome Taxonomy Database (GTDB) release 06-RS202 April 2021 (27, 28), even though GTDB currently contains fewer genomes. Based on the current GTDB taxonomy, both orders *Ca*. Troglogloeales and *Nitrospirales* are placed within the class *Nitrospiria*, but this is incongruent with analyses of their 16S rRNA phylogeny (Figure 2b). Numerous *Nitrospirota* MAGs fall outside of the three known groups of *Nitrospirota* (*Nitrospirales, Leptosprillia* and *Thermodesulfovibriona*) and are over-represented in subsurface and aquatic environments. However, 16S rRNA gene surveys indicate that members of many of the uncultivated clades exist from marine, soil and sediment environments, but are not as of yet represented by genomes (Figure 2b). Overall, while the taxonomic relationship between orders *Ca*. Troglogloeales and *Nitrospirales* and the assignment of classes in *Nitrospirota* remains to be resolved, our proposals of the genus *Ca*. Manganitrophus, family *Ca*. Manganitrophaceae, and order *Ca*. Troglogloeales are supported by both 16S rRNA gene and genome phylogenetic approaches, and additionally reveal members of a novel marine genus that possibly oxidize Mn lithotrophically.

### Genome comparison streamlines the hypothesized genes for Mn-oxidizing lithotrophy

We next compared the MAGs of members of the family *Ca*. Manganitrophaceae to understand which genes might be candidates as essential for Mn oxidation, and whether these are found in representatives of the marine genus or other members in the phylum. Four routes for Mn oxidation and electron uptake had been previously hypothesized in strain Mn1, including a Cyc2 and three different porin-dodecaheme cytochrome *c* (PCC) complexes (11). Cyc2 homologs are not only identified in the majority of *Ca*. Troglogloeales (Figure 3a), but also in other members of the phylum, including characterized clades such as acidophilic, iron-oxidizing *Leptospirilla* and nitrite or ammonia-oxidizing *Nitrospirales* (29, 30). Of the 3 PCCs in strain Mn1, only PCC_1 was found in the strains SA1 and SB1 (Figure 3a). PCC_1 was also identified in other MAGs in both marine and freshwater genera of *Ca*. Manganitrophaceae, but not in the extant MAGs and genomes of *Nitrospirota* species falling out outside of this family. These results point to PCC_1, possibly together with Cyc2, as being central to chemolithotrophic Mn oxidation by *Ca*. Manganitrophaceae.

**Figure 3.**
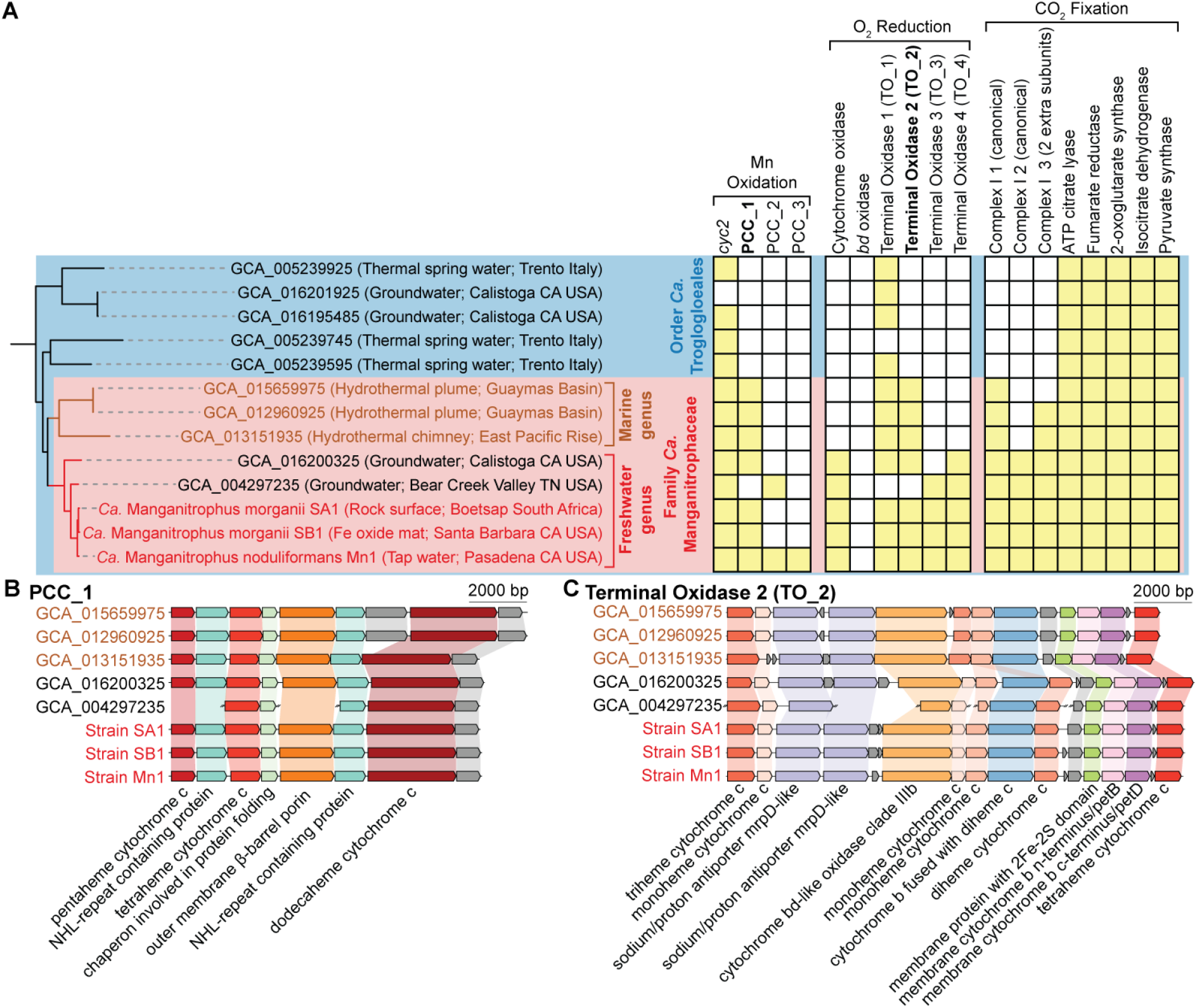
Metabolic genes and gene clusters of interest in metagenome-assembled genomes representing the order *Candidatus* Troglogloeales. **A**, The multilocus protein phylogram and the presence (yellow filled square) or absence (empty square) of genes and gene clusters of interest in the corresponding genomes. Putative functional assignments are proposed above the gene and gene cluster names. The phylogram (left) is extracted from Figure 2. **B, C**, Comparison of gene clusters of porin cytochrome c 1 (PCC_1, **B**) and terminal oxidase 2 (TO_2, **C**), both restricted to the family *Candidatus* Manganitrophaceae. Members of the freshwater genus, *Candidatus* Manganitrophus, share similar gene arrangements which differ from those representing the candidatus marine genus (in brown).

We identified five possible routes in strain Mn1 to reduce oxygen and conserve energy using electrons from Mn(II). The canonical Complex IV (*cbb3*-type cytochrome *c* oxidase) was identified in the cultivated and uncultivated members of the freshwater genus, but not in the uncultivated members of the marine genus (Figure 3a). However, the expression of this Complex IV had been observed to be low (24th percentile) in strain Mn1, especially so for a catabolic process, and therefore may not be the primary route for oxygen respiration (11). Genes for a canonical cytochrome *bd* oxidase, which has been hypothesized to reduce oxygen in *Leptospirilla* (31), were not found in strain Mn1 or other members in the order *Ca*. Troglogloeales (Figure 3a). However, genes for a number of cytochrome *bd* oxidase-like (*bd*-like) proteins that were phylogenetically distinct and predicted to have many more transmembrane helices than cytochrome *bd* oxidase (32), were identified in strain Mn1 (11). These *bd*-like oxidases are clustered with other genes potentially involved in electron transfer and energy conservation; we refer to these *bd*-like oxidase containing gene clusters as terminal oxidase (TO) complexes. While all 4 TO complexes were found in other members of *Ca*. Troglogloeales, their taxonomic distributions differed (Figure 3a). TO_1 was found in the majority of *Ca*. Troglogloeales (Figure 3a), and have been well discussed in other *Nitrospirota* including *Nitrospirales* (32, 33). TO_1 is composed of a *bd*-like oxidase clade I protein, two cytochrome *c* and a periplasmic cytochrome *b*, and was the highest expressed TO complex (98th percentile) in strain Mn1 (11). Contrasting with the more widespread distribution of the TO_1 complex across the phylum, complexes TO_2, TO_3 and TO_4 were restricted to *Ca*. Manganitrophaceae, with the latter two limited to the freshwater genus (Figure 3a). While both TO_3 and TO_4 contain two *bd*-like oxidase clade V proteins, their predicted interactions with the quinol pool differ: TO_3 encodes for an Alternative Complex III, whereas TO_4 encodes for a more canonical Complex III (11). TO_3 and TO_4 were observed to be moderately expressed at 55th and 67th percentile in strain Mn1, respectively (11). Importantly, TO_2 stands out as it was found in the majority of *Ca*. Manganitrophaceae, but as yet to be identified in any genomes outside of this family (Figure 3a). The TO_2 gene arrangement differed slightly between the two genera of *Ca*. Manganitrophaceae, but gene content was similar (Figure 3c). The TO_2 complex is composed of a membrane cytochrome *b* (similar to the petB/D or cytochrome *bf* complex) and potentially interacts with the quinone pool, a periplasmic cytochrome *b* to receive electrons in the periplasm, *bd*-like oxidase to reduce oxygen, multiple cytochrome *c* to transfer electrons, and two ion-pumping mrpD-like subunits that might be coupled to the generation or dissipation of a motive force (Figure 3c). Genes for the TO_2 complex had also been observed to be highly expressed in strain Mn1 (79th percentile) (11). Taken together, our comparative genomic analyses point to TO_2, possibly together with TO_1, as being central to Mn(II)-oxidation-dependent oxygen respiration by *Ca*. Manganitrophaceae.

### Autotrophic pathway predicted in Mn-oxidizing *Nitrospirota*

In addition to coupling the oxidation of Mn(II) to oxygen reduction, strain Mn1 was also shown to be capable of CO_2_ fixation and autotrophic growth using Mn(II) as its electron donor (11). Carbon fixation pathways such as the reverse tricarboxylic acid (rTCA) cycle, implicated in autotrophy by strain Mn1, require low potential electrons in the form of both NAD(P)H and ferredoxin (E°′ = −320 to -398 mV (21)) (34). Yet, electrons derived from Mn(II) are likely high potential (E°′ = +466 mV) (11, 12). Run in reverse, Complex I has been shown or postulated to couple the dissipation of motive force to the generation of low potential electrons and production of NAD(P)H or possibly ferredoxin (11, 35, 36).

Remarkably, in strain Mn1, 3 different Complex I gene clusters were previously identified (11). Complex_I_1 and Complex_I_2 are similar to canonical Complex I with gene clusters containing *nuoA-N* genes in order (Supplementary Figure 2). Here, phylogenomic analyses revealed that Complex_I_1 was shared by all members of both genera of *Ca*. Manganitrophaceae, whereas Complex_I_2 was restricted to members of the freshwater genus (Figure 3a). Of note, Complex_I_3 appears unique in the known biological world, having two additional ion-pumping subunits (Figure 4). This highly unusual gene cluster was found in nearly all of *Ca*. Manganitrophaceae (Figure 3a) and is not apparent in any other member of the phylum. Unusual Complex I with one additional ion-pumping subunit have been previously observed in various bacterial groups including *Nitrospirales*, termed 2M Complex I given the extra *nuoM* in the gene cluster (35), and rhizobia, termed Green Complex I in which 2 *mrpD*-like subunits have replaced the standard *nuoL* (37). Sequence comparison of Complex_I_3 subunits showed that the two MrpD-like subunits were most closely related to those in rhizobia Green Complex I, while the other subunits in the gene cluster were most closely related to those in *Nitrospira* 2M Complex I (Figure 4a). Sequence alignment of MrpD2 revealed a 26 amino acid insertion in the C-terminal amphipathic helix (HL) in all of *Ca*. Manganitrophaceae as compared to the MrpD subunits found in the rhizobia Green Complex I (Figure 4b). This type of insertion was previously identified in all gene clusters containing a second ion-pumping subunit (35). Such insertions were not unique to Complex_I_3, as they were also found in the NuoL of Complex_I_1 and Complex_I_2 in *Ca*. Manganitrophaceae (Supplementary Figure 2) and could represent evolutionary intermediates en route to being able to support additional ion-pumping subunits in the protein complex (Figure 4c).

**Figure 4.**
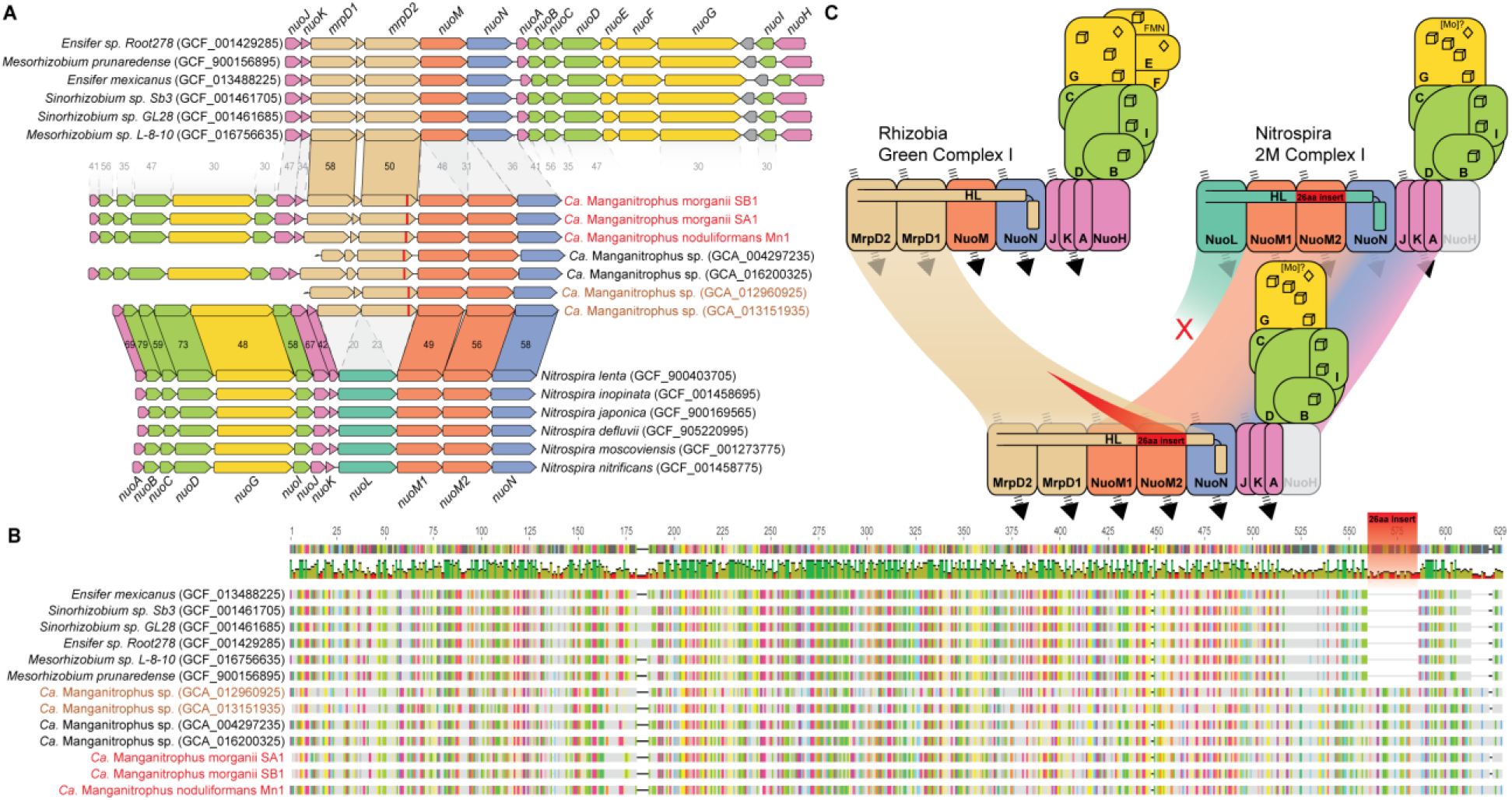
Highly unusual Complex I (Complex_I_3) with two extra pumping subunits unique to *Candidatus* Manganitrophaceae. **A**, Comparison of gene clusters of unusual Complex I with extra pumping subunits in *Ca*. Manganitrophus (middle) with their closest homologs in rhizobia (top) and *Nitrospira* (bottom). Homologs are connected between the 3 different organism clades, with values representing the average amino acid identities of proteins between the clades. NCBI accession numbers for the genome assemblies are included in brackets in the organism names. **B**, Sequence alignment of MrpD2 in Complex_I_3 in *Ca*. Manganitrophaceae reveals a 26 amino acid insert (red), compared to their closest homologs in rhizobia. **C**, Sequence comparisons reveal that Complex_I_3 in *Ca*. Manganitrophaceae is likely a hybrid between the Green Complex I in rhizobia and the 2M Complex I in *Nitrospira*. Given their sequence similarities, the two MrpD’s in Complex_I_3 could be derived from rhizobia, whereas the other components in Complex_I_3 could be derived from *Nitrospira*. The 1-2 extra pumping subunits in these unusual Complex I could enable translocation of a total of 5-6 protons or ions (as indicated by dashed arrows), compared to the 4 protons translocated by the canonical Complex I.

The majority of *Ca*. Troglogloeales and all of *Ca*. Manganitrophaceae analyzed had complete sets of genes for the rTCA cycle (Figure 3a). Only a minor difference was observed in the rTCA cycle gene content: members of the freshwater genus contained class II fumarate hydratase, whereas those of the marine genus contained class I fumarate hydratase. To further assimilate pyruvate, nearly all genes of the gluconeogenic pathway (Embden-Meyerhof-Parnas pathway) were observed in *Ca*. Troglogloeales. However, one key gluconeogenic pathway gene, namely fructose-biphosphate aldolase, was absent in strain Mn1 (11) and also appears absent from the majority of *Ca*. Troglogloeales, save for except two of the MAGs (NCBI assembly accession: GCA_004297235 and GCA_013151935). Moreover, our comparative analysis revealed that only 1 of the 5 pyruvate dehydrogenases encoded by the genome of strain Mn1 (IMG gene ID: Ga0306812_1021045-Ga0306812_1021047, Ga0306812_102629) was common to the other members of the *Ca*. Manganitrophaceae. Overall, despite apparently minor differences between their MAGs, the majority of *Ca*. Manganitrophaceae shared the same unique Complex_I_3, pathways for CO_2_ fixation and pathways for central metabolism as had been previously identified in strain Mn1.

### Core genome of *Ca*. Manganitrophaceae in marine and freshwater environments

De novo gene clustering revealed that 8 analyzed members of *Ca*. Manganitrophaceae shared a total of 895 gene clusters, which included the above-mentioned Cyc2, PCC_1, TO_1, TO_2, Complex_I_1 and Complex_I_3 (Supplementary Table 6). Several other shared genes and pathways appear noteworthy: assimilatory sulfate reduction (*sat, aprA/B, aSir*), cytochrome c biogenesis, heme exporters, 2 multicopper oxidases, and type IV pilus assembly. These confirm the basis for the ability of the cultivated strains to use sulfate as an anabolic sulfur source, make cytochrome c for anabolism and catabolism, and suggest the potential for surface twitching motility. Notably missing among the shared genes were those for the carbon-monoxide dehydrogenase complex that had been observed to be highly expressed (95th percentile) during Mn(II) dependent growth by strain Mn1 (11). Together, our comparative genomic analyses shed light on common gene sets of Mn-oxidizing chemolithoautotrophs in both marine and freshwater environments.

## Discussion

Cultivation of novel microorganisms with previously undemonstrated physiologies remains a key cornerstone to our expanding understanding of the metabolic potential of the as yet largely uncultured microbial diversity in nature (38, 39). Aerobic, Mn(II)-oxidizing chemolithoautotrophs were long theorized, but only recently demonstrated to exist in vitro in a bacterial co-culture (11). The majority member was a distinct member of the phylum *Nitrospirota, Ca*. Manganitrophus noduliformans strain Mn1, and only distantly related to any other cultivated biota (11). Curiously, the initial enrichment of Mn(II)-oxidizing chemolithoautotrophs from Caltech’s campus tap was unintentional (11). Here cultivation attempts were intentionally initiated with the specific goal of successfully establishing new Mn-oxidizing enrichment cultures. These attempts were successful using a media formulation refined during the course of the earlier study using inocula obtained from two different continents and hemispheres. Community analyses on these two new enrichment cultures revealed that the most abundant microorganisms in each were closely related to, but of a different species than *Ca*. M. noduliformans strain Mn1. The enrichment cultures also harbored a diversity of taxa varying in their relative abundances and identities (Figure 1). The results support the notion that members of the genus *Ca*. Manganitrophus are playing a key if not the central role in chemolithoautotrophic Mn(II) oxidation in the laboratory cultures examined. The results also suggest that *Ca*. Manganitrophus may not require an obligate partnership with *R. lithotrophicus* (the second species present in the previously described co-culture (11)), leaving open the possibility that its eventual clonal isolation may be possible. The phylogenomic analyses here also predict an assemblage of a marine genus within the family *Ca*. Manganitrophaceae that may also carry out this mode of chemolithoautotrophy (Figure 2, 3, and 4). However, our analyses do not exclude other members in *Nitrospirota* carrying out Mn(II) lithotrophy using a different mechanism than that we hypothesized for *Ca*. Manganitrophacae. With the increasing evidence that the *Ca*. Manganitrophaceae are distributed globally across marine and freshwater biomes (Figure 5a), taken together the reported prevalence of Mn and Mn-reducing microorganisms in the environment (14, 40), chemolithoautotrophic Mn oxidation becomes particularly important to reaching a better understanding of the redox biogeochemical cycle for manganese.

**Figure 5.**
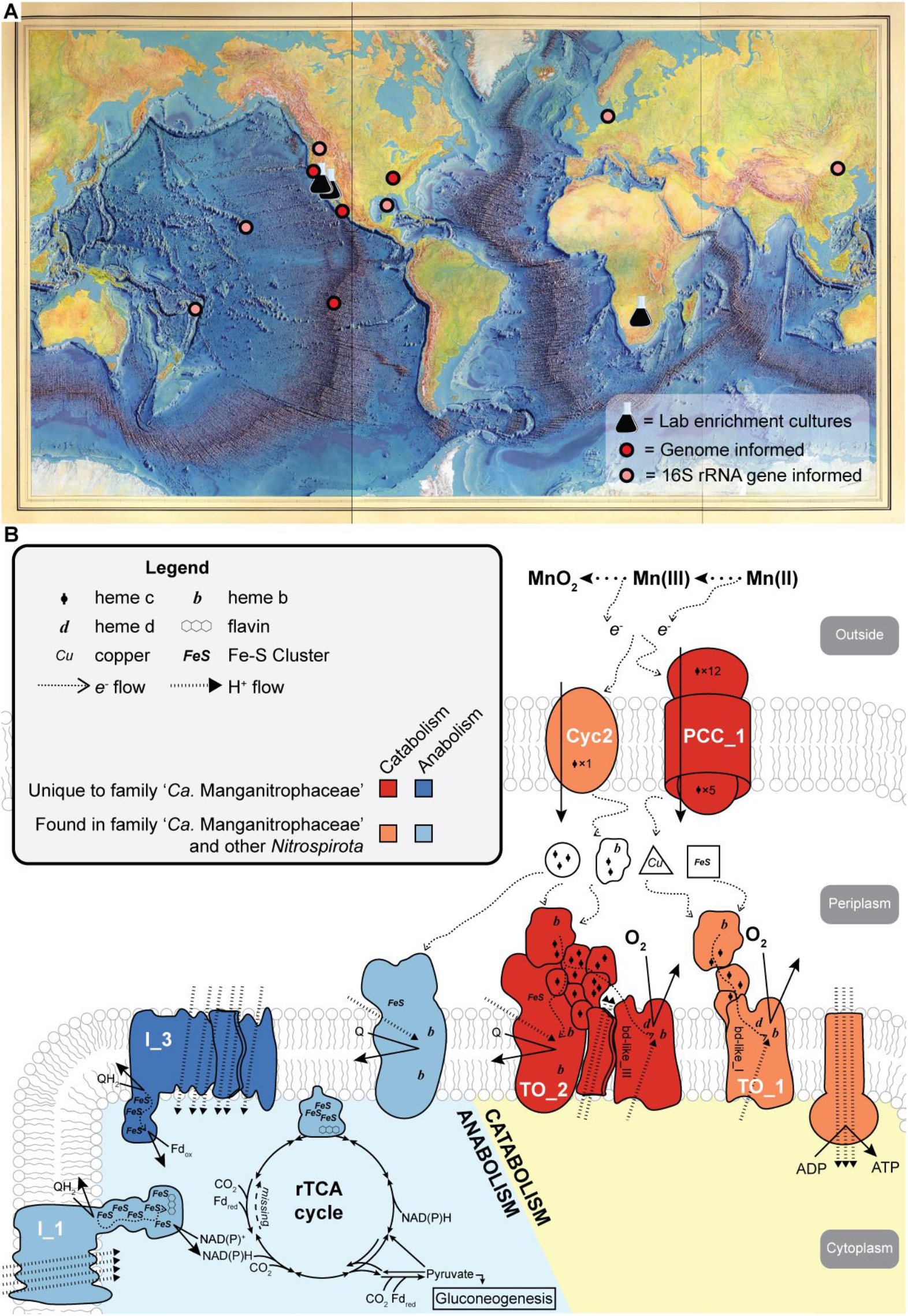
Global distribution of *Candidatus* Manganitrophaceae and key proteins and complexes putatively facilitating manganese metabolism. **A**, Distribution of cultures, metagenome-assembled genomes, and phylotypes representing *Ca*. Manganitrophaceae implies their worldwide reach in freshwater and marine environments. **B**, Cell diagram shows proposed proteins and complexes of interest to manganese chemolithoautotrophy, as deduced from representative genomes.

By comparing metagenome-assembled genomes of the 3 cultivated *Ca*. Manganitrophus strains and related but uncultivated organisms available in public genome databases, our results narrow down the list of genes in *Ca*. Manganitrophaceae that may underlie Mn(II) oxidation driven chemolithoautotrophy. Unique to *Ca*. Manganitrophaceae among all *Nitrospirota*, and perhaps across all of the biological world that has yet been analyzed, were PCC_1, as a candidate for being the initial electron acceptor during Mn oxidation; TO_2, as a candidate respiratory complex for productively coupling the electrons from Mn(II) oxidation to oxygen reduction and energy conservation (Figure 3 and Figure 5b); and Complex_I_3, as a candidate complex catalyzing reverse electron transport to generate low-potential reducing power from quinones during carbon fixation (Figure 4, Figure 5b).

While not unique to *Ca*. Manganitrophaceae, the identification of Cyc2 and TO_1 in the majority of the family members (Figure 3a and 4b), together with their comparable or even higher expression than that of PCC_1 and TO_2, respectively, in strain Mn1 (11), suggests that these two complexes may also be likely involved in Mn lithotrophy. Cyc2 is a fused cytochrome-porin protein with a single heme *c*, whereas porin cytochrome *c* (PCC) are larger complexes composed of a beta-barrel outer membrane protein and at least one multiheme cytochrome *c* (41–43). Best understood in acidophilic and circum neutral pH Fe(II) oxidation, predicted structural differences between Cyc2 and PCC homologs, specifically the smaller porin size of Cyc2 and the inner placement of heme *c* within the porin, have been suggested as meaning that Cyc2 may only react with dissolved Fe^2+^ species (29), whereas PCC homologs might react with both soluble and insoluble forms of Fe(II). In the case of Mn(II) oxidation, the reaction is thought most likely to proceed via two sequential one-electron oxidation steps (44). In that case, Cyc2 and PCC_1 might react with different forms or oxidation states of Mn (e.g. Mn(H_2_O)_6^2+^_ vs MnCO_3_, Mn(II) vs Mn(III) complexes) that have different solubilities. In comparison, known heterotrophs that catalyze Mn(II) oxidation often employ multicopper oxidase (MCO) or heme peroxidase homologs to oxidise this metal (45–47). However, the nature of these enzymes is to couple the oxidation of Mn to the direct reduction of oxygen, without a clear path for conserving any of the potential free energy energy for use by the cell. Members of *Ca*. Manganitrophaceae encodes two MCOs each (Supplementary Table 6). It is possible that these too could be involved in the lithotrophic oxidation of Mn(II); if so, it seems likely that the MCO would transfer Mn(II)-derived electrons to a periplasmic electron carriers such as cytochrome *c*, rather than directly to molecular oxygen, to be able to conserve energy for the cell (48).

Instead of the canonical cytochrome *c* oxidase common to many aerobes, *Ca*. Manganitrophaceae likely use poorly characterized terminal oxidase (TO) complexes for oxygen respiration (Figure 5b). In strain Mn1, 4 TO complexes that contain bd-like oxidases to reduce oxygen were identified, but the genes for the other components of these complexes differed between them, and likely distinguish their cellular functions (11). TO_1 contained a CISM periplasmic cytochrome *b* that may receive electrons from the periplasm, whereas TO_3 and TO_4 contained Complex III or Alternative Complex III like components that may interact with the quinone pool (11). TO_2 stands out, not only because it was found to be unique to the *Ca*. Manganitrophaceae, but also because it contains a periplasmic and a membrane cytochrome *b* that might both receive electrons from the periplasm and engage in electron transfer with the quinone pool (Figure 3c and 4b). In theory, there may be a scenario in which TO_2 bifurcates Mn(II)-derived electrons (E°′ = +466 mV) to reduce oxygen (E°’=+818 mV, via its *bd*-like oxidase) and quinones (E°’∼+113 mV, via its membrane cytochrome *b*), while using its 2 MrpD like ion-pumping subunits, unusual components of a terminal oxidase, to dissipate a membrane motive force and drive the endergonic reduction of quinones (Figure 5b).

Unusual arrangements of Complex I involving additional ion-pumping subunits may be relevant to the process of generating low-potential reducing power from quinones (35). Our analyses of Complex_I_3 examining subunit similarities, gene clustering, and the presence of specific insertions (Figure 4a and 4b) suggest an evolutionary hybridization wherein the MrpD subunits of a rhizobia-like Green Complex I replaced the NuoL of a *Nitrospira*-like 2M Complex I, with an additional HL extension needed in MrpD2 of Complex_I_3 to accommodate the second NuoM (Figure 4c). If run in reverse, this highly unusual complex, having a total of 5 ion-pumping subunits, might serve to transfer electrons from the reduced quinone pool to a carrier having a lower reduction potential than that of NADH, such as a ferredoxin required for the rTCA cycle (Figure 5b). That is, the complex could serve to dissipate the motive force built up during Mn(II) lithotrophy, by coupling the inward flow of 6 protons or sodium ions with the endergonic reduction of a ferredoxin using a quinol (Figure 4c and 5b). The additional pumping subunit would be necessary in *Ca*. Manganitrophaceae as compared to *Nitrospira* species, which have similar reverse electron transfer requirements when using high potential electron donors such as nitrite or ammonia, but are of moderately lower potential than Mn(II) derived electrons.

Based on our phylogenomic analyses, a set of shared, unique complexes in *Ca*. Manganitrophaceae, namely PCC_1, TO_2 and Complex_I_3, become prime targets for future physiological and biochemical examination, in efforts to better understand the cellular machinery enabling Mn(II)-dependent chemolithoautotrophy. Much of our proposed routes of Mn(II) oxidation are in large part informed by our existing knowledge on Fe(II) oxidation. Fe(II) oxidizers have been found in diverse marine and freshwater environments (49, 50), as is now the case for cultivated and demonstrated, as well as uncultivated and putative Mn(II) oxidizers in *Ca*. Manganitrophaceae (Figure 5a). Taxonomically, Fe(II) oxidizers have been identified in several phyla of bacteria and archaea (49, 50) and can be acidophiles or neutrophiles, mesophiles or thermophiles, phototrophs or chemotrophs, heterotrophs or autotrophs, and aerobes or anaerobes (49, 50). If such extends to the biology of energetic Mn(II) oxidation, the results gleaned here from the cultivation and phylogenomics of *Ca*. Manganitrophaceae may be only the first glimpse into the full diversity of microorganisms capable of coupling Mn(II) oxidation to growth.

## Material and Methods

### Cultivation

The enrichment procedure and manganese carbonate media composition (using 1 mM nitrate or ammonia as the N source, as noted) were described previously (11). Unless stated otherwise, culturing was performed in 10 ml of medium in 18-mm culture tubes. Cultures were transferred (10% v/v) when laboratory prepared MnCO_3_ (light pink or tan color) were completely converted to Mn oxide (dark or black color).

The South Africa inoculum was collected in June 2017 from a rock surface near a pond by a road on an exposed outcrop of the Reivilo Formation (lat. -27.964167, long. 24.454183, elevation 1107 m) near Boetsap, Northern Cape, South Africa. The rock was coated with a black material, of a texture between slime and moss. A thin, laminated green mat was observed underlying the black material. The black material reacted to leucoberbelin blue dye, indicating the presence of manganese oxides. A mixture of the black and green material was sampled using an ethanol-sterilized spatula into a sterile 15-ml tube and stored at room temperature until inoculation. The cultures were initiated in medium with 1 mM ammonia and incubated at 28.5°C. Later, some were transferred to medium with 1 mM nitrate and/or incubated at 32°C.

The Santa Barbara inoculum was collected in November 2018 from an iron oxide mat surrounded by reeds at the outflow of a rusted iron pipe (lat. 34.417944, long. -119.741130) along the side of a road in Santa Barbara, California, USA. The iron oxide mat was fluffy with a typical dark orange color. The mat was collected in a glass jar and stored at room temperature until inoculation. The enrichment cultures were incubated at 28.5°C and later some were transferred to incubate 32°C, all in the basal MnCO_3_ medium with 1 mM nitrate. The initial enrichment was transferred 5 times to confirm Mn-oxidizing activity and refine community composition prior to community and metagenomic analysis.

### Community analysis using 16S rRNA gene amplicon sequencing

Mn oxides were harvested from stationary phase enrichment cultures: 2 ml of culture containing ca. 0.15 g of Mn oxide nodules was sampled into a 2-ml Eppendorf tube and centrifuged at 8000 × g for 3 min at room temperature. After carefully removing the supernatant by pipetting, DNA was immediately extracted from the pellets using the DNeasy PowerSoil kit (Qiagen, Valencia, CA, USA) following the manufacturer’s instructions, with the bead beating option using FastPrep FP120 (Thermo Electron Corporation, Milford, MA, USA) at setting 5.5 for 45 s instead of the 10 min vortex step. DNA concentration was quantified using Qubit dsDNA High Sensitivity Assay Kit (Thermo Fisher Scientific, Waltham, MA, USA).

For 16S rRNA gene amplicon sequencing, the V4-V5 region of the 16S rRNA gene was amplified from the DNA extracts using archaeal/bacterial primers with Illumina (San Diego, CA, USA) adapters on 5’ end (515F 5’- TCGTCGGCAGCGTCAGATGTGTATAAGAGACAGGTGYCAGCMGCCGCGGTAA-3’ and 926R 5’-GTCTCGTGGGCTCGGAGATGTGTATAAGAGACAG-CCGYCAATTYMTTTRAGTTT-3’). Duplicate PCR reactions were pooled, barcoded, purified, quantified and sequenced on Illumina’s MiSeq platform with 250 bp paired end sequencing as previously described (11). Raw reads with >1 bp mismatch to the expected barcodes were discarded, and indexes and adapters were removed using MiSeq Recorder software (Illumina).

Then, the reads were processed using QIIME2 release 2020.11 (51). Briefly, forward and reverse reads were denoised using DADA2 (52) by truncating at positions 200 and 240, respectively, leaving 28 bp overlaps. Read pairs were merged, dereplicated and chimera removed with the “pooled” setting using DADA2 (52). Taxonomic assignments for the resulting amplicon sequencing variants (ASVs) used a pre-trained naive Bayes classifier on the full-length 16S rRNA genes in SILVA 138 SSURef NR99 database (53, 54). ASVs assigned to the same level 7 taxonomy were combined, and those assigned to mitochondria, chloroplast, or without taxonomy assignments were removed using the --p-exclude mitochondria,chloroplast,”Bacteria;Other;Other;Other;Other;Other”,”Unassigned;Other;Other;Other;Other;Other” setting.

### Metagenomics

Purified genomic DNA samples (2-50 ng) were fragmented to the average size of 600 bp via use of a Qsonica Q800R sonicator (power: 20%; pulse: 15 sec on/15 sec off; sonication time: 3 min). Libraries were constructed using the NEBNext Ultra™ II DNA Library Prep Kit (New England

Biolabs, Ipswich, MA) following the manufacturer’s instructions by Novogene Corporation Inc (Sacramento, CA, USA). Briefly, fragmented DNA was end-repaired by incubating the samples with an enzyme cocktail for 30 mins at 20 °C followed by a second incubation for 30 mins at 65 °C. During end repair, the 5’ end of the DNA fragments are phosphorylated and a 3’ ‘A’ base is added through treatment with Klenow fragment (3’ to 5’ exo minus) and dATP. The protruding 3’ ‘A’ base was then used for ligation with the NEBNext Multiplex Oligos for Illumina (New England Biolabs) which have a single 3’ overhanging ‘T’ base and a hairpin structure. Following ligation, adapters were converted to the ‘Y’ shape by treating with USER enzyme and DNA fragments were size selected using Agencourt AMPure XP beads (Beckman Coulter, Indianapolis, IN, USA) to generate fragment sizes between 500 and 700 bp. Adaptor-ligated DNA was PCR amplified with 9 to 12 cycles depending on the input amount followed by AMPure XP bead clean up. Libraries were quantified with Qubit dsDNA HS Kit (Thermo Fisher Scientific) and the size distribution was confirmed with High Sensitivity DNA Tapestation assay (Agilent Technologies, Santa Clara, CA, USA). Sequencing was performed on the HiSeq platform (Illumina) with paired 150 bp reads following manufacturer’s instructions (Novogene). Base calls were performed with RTA v1.18.64 followed by conversion to FASTQ with bcl2fastq v1.8.4 (Illumina). In addition, reads that did not pass the Illumina chastity filter as identified by the Y flag in their fastq headers were discarded.

The resulting reads were uploaded to the KBase platform (55), trimmed using Trimmomatic v0.36 (56) with default settings and adaptor clipping profile Truseq3-PE, and assembled using Spades v3.11.1 (57) with default settings for standard dataset. Manual binning and scaffolding were performed using mmgenome v0.7.179 based on differential coverage and GC content of different metagenomes to generate the MAG for the most abundant organism. MAGs were annotated using the Rapid Annotations using Subsystems Technology (RAST) (58–60) and NCBI Prokaryotic Genome Annotation (61) Pipelines. Average nucleotide identities and reciprocal mapping of MAGs were done using fastANI v1.32 (24). Average amino acid identities were done using enve- omics tool AAI calculator (26). De novo gene clustering was done using anvio v7 with default parameters (62). Comparison of Complex I gene clusters was done using protein-protein BLAST with default parameters (63) to the RefSeq Select proteins database (64). Alignment of Complex I gene sequences was done using MUSCLE v3.8.1551 with default parameters (65).

### Phylogenetic analyses

For genome phylogeny, 433 publicly available genome assemblies in the NCBI Assembly Database (61) fell within the phylum *Nitrospirae* (Taxonomy ID 40117) (66) and 6 publicly available genomes in the genomic catalog of Earth’s microbiomes dataset (67) fell within the phylum *Nitrospirota* under the headings Nitrospirota and Nitrospirota_A (27) and were analysed (as of March 30, 2021). For 16S rRNA gene phylogeny, 16s rRNA genes from the MAGs of Nitrospirota from the enrichment metagenomes, as well as the genome assemblies were retrieved using CheckM v1.1.2 (68) ssu_finder utility. Sequences less than 900 bp were excluded. The 16S rRNA gene sequences were aligned using SINA v1.2.11 (69) and imported into SILVA Ref Database Release 138.1 (53). 104 16S rRNA gene sequences, including 5 different outgroup sequences (*Desulfovibrio vulgaris, Ramlibacter tataouinensis TTB310, Nitrospina gracilis 3/211, Acidobacterium capsulatum, Candidatus* Methylomirabilis oxyfera), with 1508 nucleotide positions were exported with bacteria filter excluding columns with mostly gaps from the ARB software v6.0.2 (70). Bayesian phylogenetic trees were constructed using MrBayes v3.2.7 (71) with evolutionary model set to GTR + I + gamma, burn-in set to 25% and stop value set to 0.01, and edited in iTOL v6 (72). For concatenated multilocus protein phylogeny, marker proteins from 104 genomes including the same 5 outgroup species were identified and aligned using a set of 120 ubiquitous single-copy bacterial proteins in GTDB v0.2.2 (27). The protein alignment was filtered using default parameters in GTDB v0.2.2 (27) (the full alignment of 34744 columns from 120 protein markers were evenly subsampled with a maximum of 42 columns retained per protein; a column was retained only when the column was in at least 50% of the sequences, and contained at least 25% and at most 95% of one amino acid). The resulting alignment with 5040 amino acid positions was used to construct the multilocus protein phylogeny using MrBayes v3.2.7 (71) as above except the evolutionary model was set to invgamma and a mixed amino acid model.

### Data Availability

The partial 16S rRNA gene amplicon sequences of enrichment cultures, and metagenome-assembled genomes of *Candidatus* Manganitrophus morganii strains SA1 and SB1 have been deposited in National Center for Biotechnology Information (NCBI) under BioProject PRJNA776098.

## Acknowledgements

We thank Stephanie Connon for assistance with 16S rRNA gene amplicon sequencing and analysis. This work was supported by NASA Astrobiology Institute Exobiology grant #80NSSC19K0480; and by Caltech’s Center for Environmental Microbial Interactions and Division of Geological and Planetary Sciences. Fieldwork in South Africa by UFL was supported by Woodward Fischer (Caltech) under grants from the NSF (IOS-1833247) and the Packard Foundation.

